# The efficiency of Grignard Pure™ to inactivate airborne SARS-CoV-2 surrogate

**DOI:** 10.1101/2022.08.19.504307

**Authors:** Grishma Desai, Gurumurthy Ramachandran, Emanuel Goldman, Antony Galione, Altaf Lal, Toni K. Choueiri, Andre Fay, William Jordan, Donald W. Schaffner, Jack Caravanos, Etienne Grignard, Gediminas Mainelis

## Abstract

Grignard Pure™ (GP) is a unique and proprietary blend of Triethylene Glycol (TEG) and inert ingredients designed for continuous antimicrobial treatment of air. GP received approval from the US EPA under its Section 18 Public Health Emergency Exemption program for use in seven states. This study characterizes the efficacy of GP for inactivating MS2 bacteriophage – a non-enveloped virus widely used as a surrogate for SARs-CoV-2. Experiments measured the decrease in the airborne viable MS2 concentration in the presence of different concentrations of GP from 60 to 90 minutes, accounting for both natural die-off and settling of MS2. Experiments were conducted both by introducing GP aerosol into air containing MS2 and by introducing airborne MS2 into air containing GP aerosol. GP is consistently able to rapidly reduce viable MS2 bacteriophage concentration by 2-3 logs at GP concentrations of 0.02 mg/m^3^ to 0.5 mg/m^3^ (corresponding to TEG concentrations of 0.012 mg/m^3^ to 0.287 mg/m^3^). Related GP efficacy experiments by the US EPA, as well as GP (TEG) safety and toxicology, are also discussed.

**Synopsis:** Limited research on the germicidal properties of triethylene glycol against airborne pathogens was conducted during the 1940s and 50s. This paper investigates the inactivation rate of airborne bacteriophage MS2 by Grignard Pure^™^ product, containing a unique and proprietary blend of Triethylene Glycol (TEG) and inert ingredients.

## A. Introduction

The COVID-19 pandemic has raised awareness of the airborne transmission of infectious diseases^1^, including transmission by humans. When infected individuals speak, cough, sneeze, or sing, they release both large respiratory droplets and smaller airborne microdroplets or aerosol particles (< ~ 5 μm)^2^. Large droplets quickly settle on surfaces within 6-10 feet of the source due to gravity, while smaller aerosol particles (usually <5 μm) can stay afloat for minutes and even hours, especially if aided by air currents^3^. Laboratory studies have shown the presence of infectious SARS-CoV-2 in such human-generated microdroplets, and SARS-CoV-2 can remain viable for up to 16 hours, with a half-life for the viability of 0.5 – 3.3 hours depending on the size distribution of the respiratory aerosol^4^. Respiratory droplets with a diameter of 0.09 μm containing one virion per droplet may persist for hundreds of hours, whereas 0.4 μm respiratory droplets containing SARS-CoV-2 virus may remain infectious for only a few hours^4^. Particles containing the virus have to be either removed from the air or the virus in those particles has to be inactivated to reduce the risk of exposure to airborne viable viral particles.

Many products are available and approved for disinfecting hard and soft surfaces harboring SARS-CoV-2 (e.g., US EPA’s list N ^5^). However, there is a clear need for technologies that can reduce the airborne transmission of the SARS-CoV-2. Among the potential substances to inactivate airborne biological agents, triethylene glycol (TEG) was demonstrated to have germicidal properties more than 70 years ago^6^. Aerosolized TEG is almost 100 times more potent against respiratory pathogens compared to TEG in liquid form^6^. Puck demonstrated in the 1940s that the lethal effect of TEG occurs once a sufficient amount of TEG vapor molecules condenses on particles containing the microbes^7^. Robertson, et al. (1943) confirmed that TEG vapor was an effective decontaminant for airborne infectious agents, including viruses causing influenza, meningopneumonitis, and psittacosis^8^. Bacteria found to be susceptible to TEG vapor include pneumococci type I, II and III, beta hemolytic streptococci group A and C, staphylococci, influenza bacilli, *Escherichia coli*, and *Bacillus aerogenes*^6^. As little as 2 – 5 mg/m^3^ of TEG in the air was sufficient to produce “maximum germicidal action” against various airborne infectious agents^6^.

Pure TEG is difficult to safely aerosolize for air treatment purposes due to fire risk^9^. Grignard Pure™ was developed using TEG as the active ingredient and contains water and propylene glycol ingredients to aid in faster evaporation while preventing fire hazards^10^. Since the active ingredient in Grignard Pure™ is TEG, we hypothesized that Grignard Pure™ has the potential to act as an antimicrobial agent. The goal of this paper is to investigate the efficacy and potential application of Grignard Pure™ as an airborne antimicrobial agent that can provide a much-needed additional layer of protection for indoor spaces.

## B. Materials and Methods

### Aerosolization of Grignard Pure™

Grignard Pure ™ (GP) includes TEG as the active ingredient and propylene glycol and deionized water as described in WO 2021/226232^11^. GP was utilized in its undiluted form and aerosolized through proprietary vaporization or nebulization devices. Vaporizers pass the GP solution over a heating block, where GP is heated above its boiling point and vaporized. The vapor released in the target air space (e.g., test chamber) rapidly condenses to form fine droplets producing a visible aerosol, i.e., haze or fog. Vaporizing dispersion devices can be handheld or free-standing. The Nimbus handheld vaporizing device (Grignard Pure LLC, Rahway, NJ) was used in these studies to treat the test chamber with a single, four-second release of GP. Two stand-alone vaporizing units, the Clearify and the Amhaze (both from Grignard Pure LLC), were used to treat the chamber air with a controlled time release of GP, where GP is periodically injected to maintain a set concentration. Nebulizers aerosolize the GP solution and disperse it through a fine-tipped nozzle. The Aura stand-alone nebulizing device (Grignard Pure, LLC) was used to treat the chamber air in a controlled time release mode. The target GP aerosol concentration in a chamber was achieved by adjusting the output volume and the duty cycle of the employed devices. The resultant GP aerosol concentration was measured as described below and used to determine the total airborne TEG concentration.

### Measurement of Triethylene Glycol (TEG) and Grignard Pure™ concentrations in the air

The mass concentration of Grignard Pure™ aerosol was correlated to the total concentration of TEG in the air (aerosol and vapor). These experiments and the resulting correlation curve are described in Supporting Information (Figures S1 and S2, and Table S1).

### Testing of TEG Efficacy Against Airborne Virus

Two different laboratories referred to as Lab 1 and Lab 2 below, performed studies to investigate TEG aerosol efficacy against airborne MS2 bacteriophage. Experimental setups were slightly different and are described below. Total TEG concentration in the air was determined based on GP aerosol concentrations as described in Supporting Information.

### Aerosol Test Chambers

All testing in Lab 1 was performed in a negative pressure aerosol chamber measuring 3 m (H) × 3 m (W) × 2.4 m (D) made from polycarbonate plastic with a thickness of 0.038m (Figure S3 in Supporting Information). The chamber, including the walls, glove ports, and sampling ports, was thoroughly cleaned with 1:100 diluted household bleach solution before initiating the test and between experiments. On each test day, prior to each experiment, the sampling ports were wiped with 1:100 household bleach solution.

All testing in Lab 2 was conducted in a fully enclosed, 2.7m (H) × 2.7m (W) × 2.1(D) 304 stainless steel sealed chamber equipped with various sampling ports (Figure S4 in Supporting Information). Following each test, the chamber was evacuated and purged with HEPA-filtered air for a minimum of 20 mins. The chamber surfaces were wiped with a 50/50 mixture of 95% isopropyl alcohol as well as DI water. Thirty percent hydrogen peroxide was nebulized into the chamber for 20 mins between trials.

### Inoculum Preparation and Aerosolization

MS2 bacteriophage (ATCC 15597-B1) and the host microorganism *Escherichia coli* (ATCC 15597) were used in all experiments. MS2 is a small non-enveloped virus and has been used as a surrogate for the more sensitive enveloped SARS-CoV-2. In addition, MS2 is well characterized and has frequently been used as a surrogate for other pathogenic viruses (e.g., influenza virus and SARS-CoV-1) in aerosolization and inactivation studies^12^.

Prepared viral stocks in Lab 1 were stored at −70°C ± 10°C until ready to be used for testing. On the day of testing, frozen stocks were removed from the refrigerator and allowed to thaw at room temperature. Host culture was grown in 10 ml of tryptic soy broth (Hardy Diagnostics, Santa Maria, CA) at 36 ±1°C for 6 – 24 hours. The inoculum was prepared by diluting viral stock in phosphate buffered saline solution to a target concentration of ≥ 1.0 × 10^7^ PFU/ml. Concentration was determined by performing a standard plate count by enumerating the inoculum prior to nebulization. MS2 was introduced into the chamber using two Collison 6-jet nebulizers (CH Technologies, Westwood, NJ) operated at 10 psi pressure. Nebulizers were prepared by adding 15 – 20 ml of inoculum inside a biological safety cabinet (NuAire, Plymouth, MN). A 6-inch desk clip fan was used to mix the air inside the chamber.

Working stock cultures in Lab 2 were prepared using aseptic techniques in a Class 2 biological safety cabinet (Labconco, Fort Scott, KS), following standard preparation methodologies. Approximately 250 mL of *E. coli* was prepared in tryptic soy broth and incubated for 24 – 48 hours with oxygen infusion (1 cm^3^/min) at 37°C. Bacterial stock concentrations were ~10^9^ CFU/mL as determined through triplicate plating and enumeration. The bacterial suspension was infected during the logarithmic growth cycle with MS2 bacteriophage. After 18 hours of incubation, cells were lysed, and the cellular debris discharged by centrifugation. MS2 bacteriophage stock yields were calculated through small drop plaque assay plating and enumeration and determined to be greater than 10^11^ (PFU/mL) with a single amplification procedure. The virus was introduced into the chamber through a 24-jet Collison nebulizer (CH Technologies, Westwood, NJ). The Collison nebulizer was filled with 40ml of inoculum and operated at 40 psi. This lab used AM520 aerosol photometer (TSI, Inc.) to determine Grignard Pure™ aerosol mass concentration. This device was calibrated using Arizona Road Dust as described in Supporting Information.

### Grignard Pure™ Dispersion Device Preparation

GP was used in its undiluted form in all experiments. Dispersion devices were primed outside the test chambers for about 5 mins at predetermined operational settings prior to each test

### Aerosol Efficacy Chamber Runs

Efficacy studies at both labs were conducted at six aerosol concentrations of GP ranging from 0.02 mg/m^3^ to 0.5 mg/m^3^ with corresponding total TEG concentrations of 0.012 mg/m^3^ to 0.287 mg/m^3^. Different GP (and corresponding TEG) concentrations were used to investigate whether the selected concentration range affects the efficacy of GP against MS2.

### Test Scenarios

Two different contact protocols were used: 1) single shot-burst (4 sec) release of GP into the MS2 aerosol and 2) a controlled time release of GP to maintain a set GP concentration. Both test scenarios are illustrated in Figure 1.

**Figure 1.**
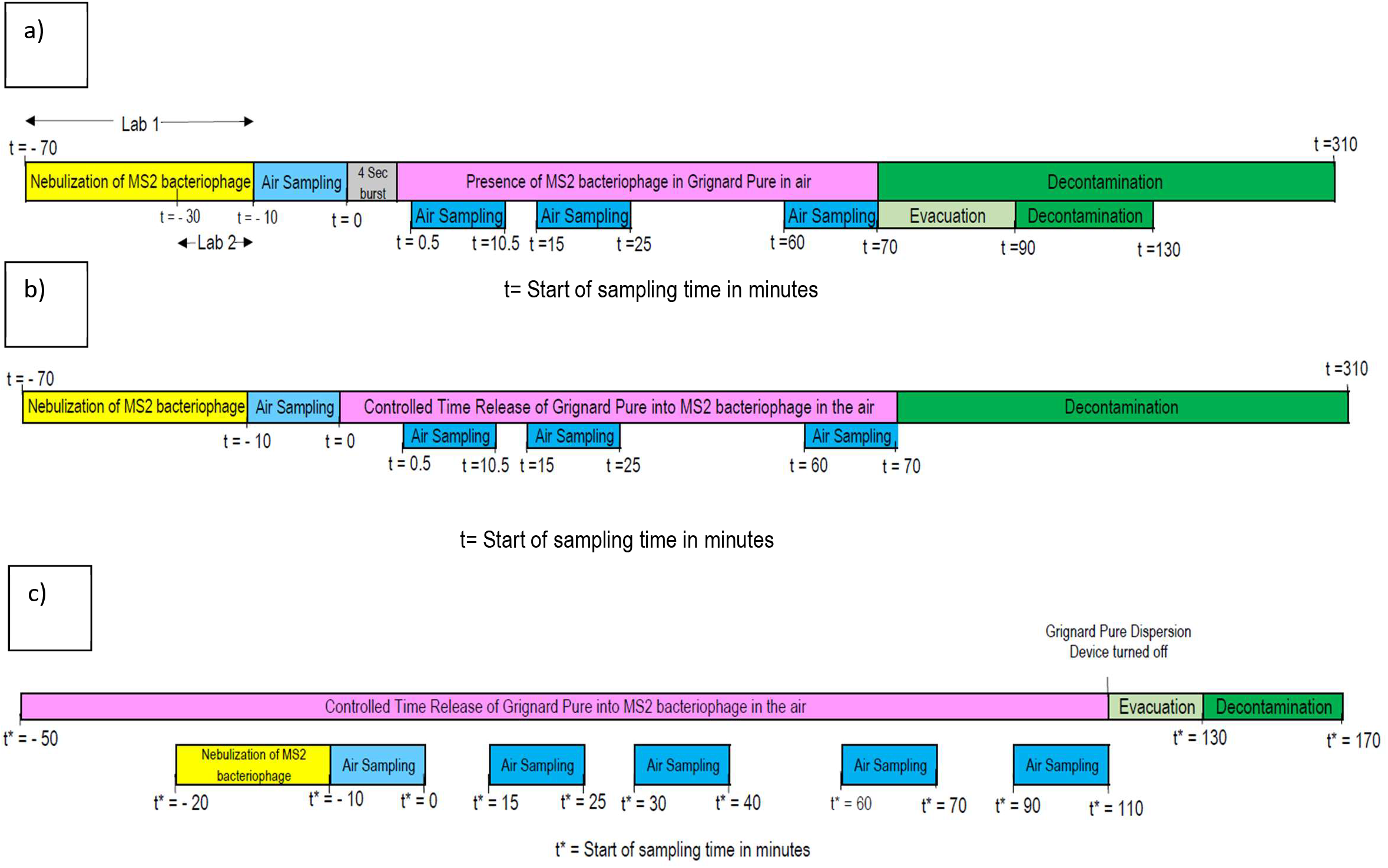
Timeline of experiments to investigate the efficacy of Grignard Pure™ (GP) against MS2 bacteriophage: a) timeline for 4-second burst testing of GP efficacy at Lab 1 and Lab 2. GP was released using Nimbus vaporizing device; b) timeline for controlled time release testing of GP efficacy in Lab 1. GP was released using Amhaze; c) timeline for controlled time release testing of GP efficacy in Lab 2. GP was released using either the Clearify or Aura. Here, MS2 was nebulized into airborne GP, and thus, to differentiate the time zero in these experiments from experiments in part b, time is denoted as “t*.”

The sequence of events for the first scenario is shown in Figure 1a. MS2 bacteriophage was aerosolized for 60 min by Lab 1 or 20 min by Lab 2, and then a sample of airborne MS2 was collected at t = −10 min to determine the airborne MS2 concentration before the air treatment. Once this sample was collected, a single, four-second burst of GP was introduced into the chamber using the Nimbus vaporizing device (t=0 min). The airborne MS2 samples were then collected at starting time points t= 0.5, 15, and 60 min. Once the last sample was collected, the test chambers were evacuated/decontaminated. During the control experiments (i.e., no GP was introduced into the chamber air), samples of airborne MS2 were collected at the same time points of t= −10, 0.5, 15, and 60 min.

In the second scenario, the two labs used slightly different protocols (Figs. 1b and 1c). Lab 1 (Figure 1b) started nebulizing the MS2 phage at t = − 70 min for 60 minutes ± 30 seconds to reach the target concentration of the test organism. Then the nebulization stopped, and the airborne virus was collected for 10 min, from t = −10 min to 0 min. Immediately after the sample was collected, a controlled time release of GP was initiated, and it lasted from t = 0 until t = 70 min. During this time, three 10-min samples of airborne MS2 were collected with sampling start times of t = 0.5, 15, and 60 min. At the end of the last sample, the GP aerosolization was stopped, and the chamber was decontaminated. A series of controlled time release experiments were performed at Lab 1 at several different airborne TEG concentrations: 0.063, 0.186, 0.235, and 0.287 mg/m^3^. (The corresponding GP concentrations were 0.1, 0.32, 0.4, and 0.5 mg/m^3^). The experiments with 0.235 mg/m^3^ concentration were performed in triplicate, while experiments with other concentrations were single experiments, adding to the totality of evidence about the treatment efficacy of GP. Experiments at each GP concentration had their own separate control experiments. The GP was aerosolized using the Amhaze, a stand-alone vaporizing device, and it periodically vaporized GP to maintain its concentration in the air. During the control experiments (i.e., GP was not introduced into the chamber air), samples of airborne MS2 were collected at the same time points of t= −10, 0.5, 15, and 60 min.

Lab 2’s scenario for the controlled time release was different from that of Lab 1 in the initial phases of the experiments (Figure 1c) to reflect the testing protocol used by the US EPA, which tested different air treatment technologies, including GP^13^. During these experiments in Lab 2, GP was released into the air first and continued to be periodically released for the duration of the experiment to maintain its steady concentration. Twenty minutes after the release of GP was started, the aerosolization of MS2 was initiated for 20 minutes. After 20 minutes of nebulization, it was assumed that the MS2 was evenly distributed inside the chamber, and the first 10-min sample of airborne MS was collected. The sample starting point was marked as time zero. In order to differentiate time points in these experiments from the time points described above, the “t*” symbol will be used. The subsequent samples of airborne MS2 were collected at t*= 15, 30, 60, and 90 min (again, these time points correspond to the EPA’s testing protocol). After the last sample, GP aerosolization was stopped, and the chamber was evacuated and decontaminated. These experiments in Lab 2 were performed with the Clearify, a stand-alone vaporizing device, and the Aura, a stand-alone nebulizing device. For the Clearify, target GP aerosol concentration was 0.16 mg/m^3^ equating to a total TEG concentration of 0.092mg/m^3^. For the Aura, the target GP aerosol concentration was 0.04 mg/m^3^ equating to a total TEG concentration of 0.025 mg/m^3^. During the control experiments (i.e., GP was not introduced into the chamber air), samples of airborne MS2 were collected at the same time points of t*= 0, 15, 30, 60, and 90 min.

### Collection of airborne MS2 and analysis procedure

Lab 1 used Biosamplers (SKC Inc., Eighty-Four, PA) operated at 12.5 L/min and filled with 20 ml of sampling media composed of phosphate buffered saline with 0.1% Tween 80 to collect air samples. Spatially distributed triplicate samples were collected for 10 minutes ± 10 seconds at each sampling time point. After sampling, the samplers were moved to a biosafety cabinet for processing. Each sampler’s neck was rinsed with 5 ml of sterile phosphate buffered saline and allowed to drain into the collection cup of the vessel. The liquid was transferred into a sterile 50 ml conical vessel, and the total liquid volume was observed and recorded. Samples were diluted in a series of ten-fold dilutions in phosphate buffered saline to observe a countable range of plaques, i.e., 25 – 250 colonies per plate. The dilutions were plated using the pour plate technique. The dilutions of MS2 samples were plated on 50% tryptic soy agar (TSA), which was supplemented with 0.100 ml per plate with *E. coli* (ATCC 15597). Plates were swirled and then allowed to solidify prior to incubation. The plates and controls were incubated at 36°C ± 1°C for 18 – 24 hrs. The number of plaque forming units (PFU) in each sample was enumerated and converted to airborne concentration (PFU/m^3^). For dilution series with counts < 25 on the least diluted plates, counts less than 25 were used in calculations. For dilution series with no counts, the limit of detection was used to estimate virus concentrations. The limit of detection of the MS2 for the bioaerosol concentrations was 80 PFU/m^3^

In Lab 2, two AGI-30 impingers (Ace Glass Inc., Vineland, NJ) located at opposite corners of the chamber were used to collect air samples at their nominal flow rate of 12.5 L/min. The impingers contained 20 ml of sampling media composed of sterilized phosphate buffered saline with 0.005% v/v Tween 80. Aliquots of impinger samples were plated in triplicate on tryptic soy agar media using the small drop plaque assay technique over a minimum 3-log dilution range. The samples were diluted in the phosphate buffered saline solution using a 1:99 dilution of overnight MS2 bacteriophage plating stock *E. coli* (ATCC #15597). Plates were incubated for 24-48 hours, and the resulting PFUs were counted and converted to PFU/m^3^. The limit of detection of the MS2 for the bioaerosol concentrations was 1000 PFU/m^3^

### Calculation of Grignard Pure™ treatment efficacy

In control experiments (i.e., without Grignard Pure™ present in the air), the concentration of viable airborne MS2 bacteriophage decreases due to natural die-off and settling (NDOS); when the air is treated by Grignard Pure™, the MS2 loses viability due to natural die-off and settling and the inactivating action of Grignard Pure™. To determine the base-10 log reduction (Lg) in MS2 viability caused by Grignard Pure™ (*LR_GP_*), the log reduction in MS2 viability during control experiments (*LR_NDOS_*) must be subtracted from the log reduction during treatment experiments (*LR_NDOS+GP_*). Since the virus titers in Lab 1 during the control and treatment experiments were different, *LR_NDOS_* and *LR_NDOS+GP_* were first calculated separately relative to the sample collected starting at t = − 10 min and then used to determine (*LR_GP_*) for each sampling time *x*, as follows:

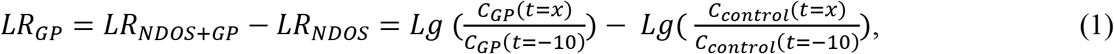

where *C_control_* is airborne viable virus concentration during control experiments (i.e., air not treated by GP), and *C_GP_* is the airborne virus concentration when the air was treated by GP. *LR*_*NDOS*+*GP*_ could also be called gross log reduction, while *LR*_*GP*_ is net of effective log_10_ reduction.

In Lab 2, the virus titers during the control and treatment experiments were the same, and Eq. 1 could be simplified to the following equation for each sampling time point *x*:

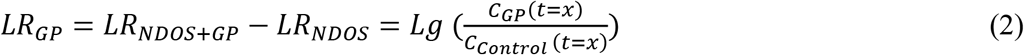

Each *LR* for each time point can be easily converted into percent reduction (*PR*):

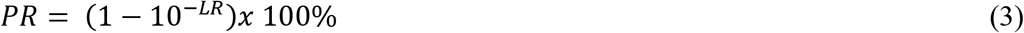

## C. Results and Discussion

### Efficacy of a four-second burst of TEG against the airborne virus

The concentrations of viable airborne MS2 bacteriophage and its reduction over time when treated with a single four-second burst of GP at Lab 1 and Lab 2 and the resulting net log reduction are shown in Table S2 in Supporting Information and Figure 2, respectively.

**Figure 2.**
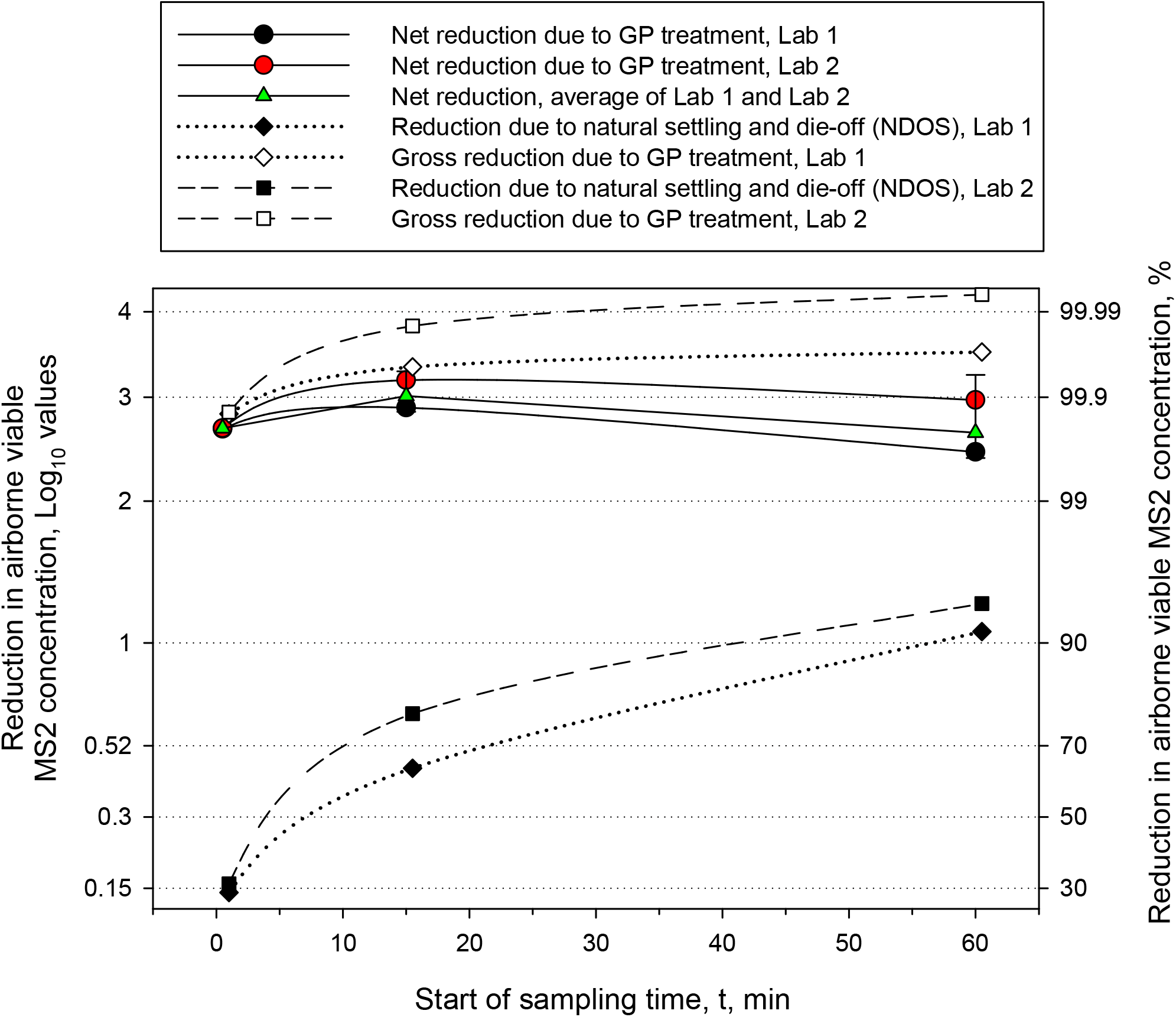
The log and percent reduction in viable airborne MS2 phage concentration at different sampling points when treated with a single 4-second burst of Grignard Pure™ aerosolizee by the Nimbus, a handheld vaporizing device. The experimental sequence in Lab 1 and Lab 2 was identical. In this experiment, MS2 was aerosolized first and then a 4-second burst of Grigard Pure™ was introduced. The airborne TEG concentration was not recorded.

Viable MS2 concentration decreased by 0.15 log (30%) at 0.5 min, and the decrease reached over 1 log (90%) at a sampling time of 60 min in all experiments due to natural die-off and settling. The gross inactivation was 3 (99.9 %) logs at 0.5 min and 4 logs (99.99%) at 15 and 60 min. This yielded the net log reduction of 2.6 in the aerosol concentration of MS2 bacteriophage when sampled at t=0.5 min after treatment by GP.

At the t= 15 min sample, Lab 1 observed a net log reduction of 2.89 in MS2 viable concentration, while the net log reduction value reported by Lab 2 was 3.18. After 60 min of treatment, both labs recorded slightly lower net reduction values compared to t = 15 min: 2.44 by Lab 1 and ~3 by Lab 2.

Data from both labs are in good agreement, which is important because the two labs used slightly different virus preparation protocols and different initial viable virus concentrations. Lab 1 started experiments with 3.4 × 10^8^ PFU/m^3^, while Lab 2 started with almost two orders higher virus concentrations, 6.7 × 10^10^ PFU/m^3^. When the data from the two labs are averaged, the net log reduction at t=0.5 min is 2.6, and at t=15 min, the net reduction is 3, i.e., 99.9% of the viable virus was eliminated. In addition, the log reduction in MS2 concentration with GP treatment is statistically significantly different (p<0.05) from the log reduction in MS2 due to natural die-off and settling for all three time points, according to a paired t-test. The observed reduced inactivation at t= 60 min is due to the limit of detection issues, i.e., too few viable viruses remained in the air after the treatment.

### Efficacy of controlled time release of Grignard Pure™ against airborne virus particles

The log reductions in airborne viable MS2 bacteriophage concentration during control and treatment experiments at different sampling time points are presented in Figure 3, while the concentration values are given in Table S3 in Supporting Information. The results show that GP is highly effective and yields 1-2.5 net log reduction in MS2 concentration in samples that were initiated just 30 s post-treatment (Fig. 3c). In samples that were collected starting 15 min after the treatment, the net log reduction ranged from 2 at the lowest tested TEG concentration of 0.063 mg/m^3^ to 3.1 at the highest tested TEG concentration of 0.287 mg/m^3^. In general, for 0.5-minute and 15-minute sampling times, the inactivation seemed to increase somewhat with increasing TEG concentration. At 60 min sampling time, compared to the previous sampling times, the net log reduction either stayed the same (TEG = 0.063 mg/m^3^), increased (TEG = 0.186 and 0.235 mg/m^3^) or slightly decreased (TEG = 0.287 mg/m^3^). The steady or decreasing inactivation at the 60 min sampling point compared to 0.5- and 15-minute sampling points could be attributed to viable virus concentration approaching the limit of detection of the sampling and analytical method. When data from all experiments are pooled together, regardless of the GP concentration, the observed log reduction in viable MS2 concentration with GP treatment is statistically significantly different (p<0.01) from the log reduction in airborne MS2 concentration due to natural die-off and settling for all three time points, according to a paired t-test.

**Figure 3.**
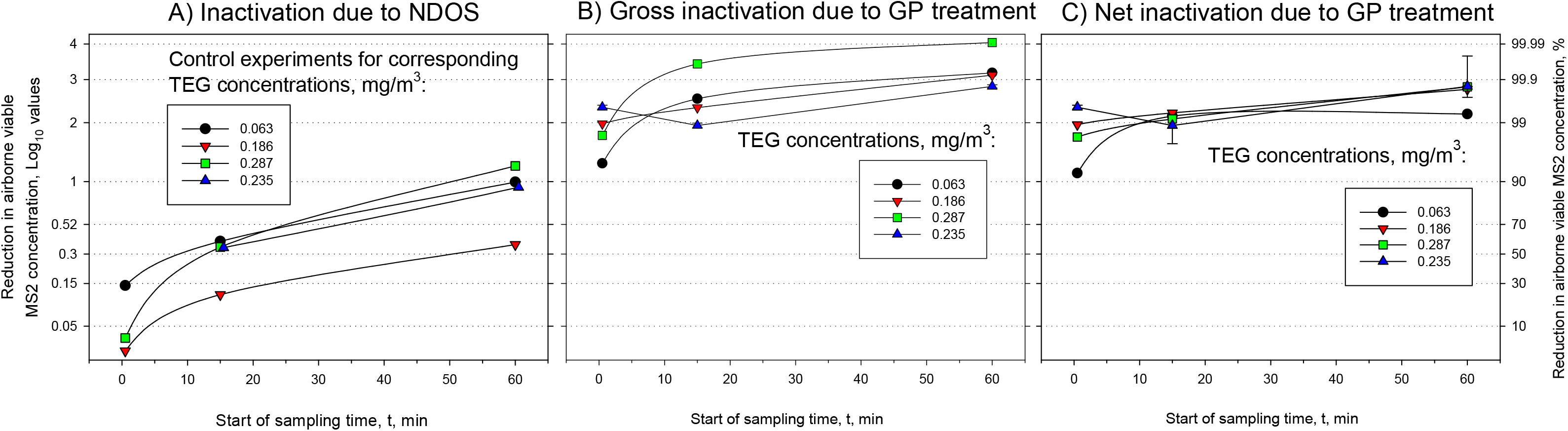
The log and percent reduction in viable MS2 concentration due to air treatment by different concentrations of Grignard Pure™ and the resulting TEG at different sampling points. In this experiment, MS2 was aerosolized first and then Grigard Pure™ was aerosolized into MS2 in a controlled time release mode using the Amhaze, a stand-alone vaporizing device.

The log reductions in airborne viable MS2 bacteriophage concentration during control and treatment experiments are presented in Figure 4. The observed concentrations of viable airborne MS2 bacteriophage are in Table S4 in Supporting Information. A gross log reduction of 2.57 in viable MS2 concentration can be seen at time zero when the GP was released using the Clearify device according to the schematic in Figure 1c; both devices yielded a 1.60 net log reduction. At the 15 min sampling time, the net log reduction increased to 3.2 with Clearify and 2.5 for Aura. As time progressed, the net log inactivation of MS2 steadily increased to approximately 3.3 when the Aura device was used. For the treatment using the Clearify device, the net log reduction remained steady above 3 due to the low remaining viable virus concentration in the air.

**Figure 4.**
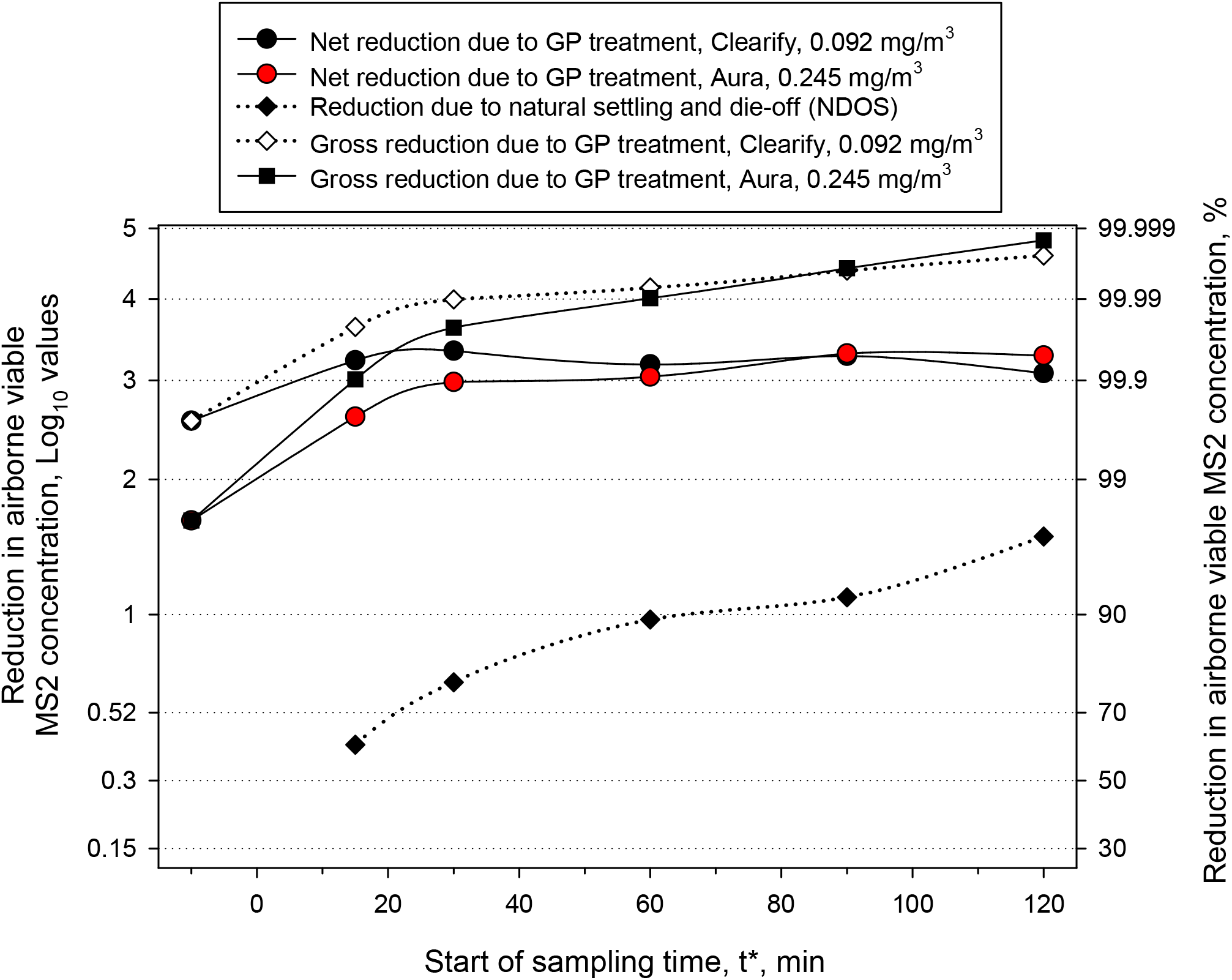
The log and percent reduction in the concentration of airborne viable MS2 bacteriophage at different sampling points when it was treated with a controlled time release of Grignard Pure™ using the Clearify Device, a stand-alone vaporizing device and the Aura, a stand-alone nebulizing device at Lab 2. In this experiment, MS2 was aerosolized into airborne Grignard Pure.

### Comparison with testing by the US EPA

In addition to the testing conducted by Grignard Pure LLC at Lab 1 and Lab 2, GP was also evaluated by the US EPA’s Office of Research and Development in May – June 2021 as part of their COVID-19 Research^14^. The main objective of the EPA’s research was to evaluate the efficacy of different types of aerosol treatment technologies in reducing airborne virus concentrations using a large-scale test chamber and a standardized testing approach^14^.

The effectiveness of GP against the MS2 bacteriophage was evaluated by the EPA in two test scenarios. In the first, the MS2 bacteriophage was first introduced into the chamber as an aerosol, the initial bioaerosol sample was taken to determine the virus concentration at time = 0 min, and then GP was added to the chamber, similar to the test scenario in Lab 1 as shown in Figure 1b^13^. This allowed for a direct assessment of the efficacy of inactivation of GP at a high concentration of the MS2 bacteriophage in the chamber air as a function of time since the product introduction^13^. In the second scenario, the GP was first added to the chamber environment at the desired concentration before the MS2 bacteriophage was aerosolized in the chamber, similar to the test scenario in Lab 2, as shown in Figure 1c^13^. This scenario more directly assessed the continued use of GP in occupied spaces, where virus could be introduced by an infected individual(s) into a space where the target concentration of GP is maintained^13^. The materials and methodology employed by the US EPA are available on the official US EPA COVID-19 Research webpage^13^.

With the first test scenario, at the sampling time of 15 min, the percentage reduction observed at the US EPA was 95.5% and 99.76% (1.3 and 2.6 logs)^13^, similar to results from Lab 1. With both tests, there was an increased percentage reduction of MS2 bacteriophage as time passed. At 60 minutes, the testing at the US EPA achieved a 97.6% (1.6 logs) reduction, and the testing at Lab 1 achieved a 99.8% (2.7 logs) reduction. These results further confirm GP’s ability to achieve at least 1.0 – 2.5 log reduction at the first sampling time. The difference in results may be attributed to the size of the chamber, inoculum preparation, and aerosol sample collection method. Data for the second test scenario showed a similar trend in reducing viable MS2 bacteriophage concentration at both the US EPA and Lab 2. At time zero, a 99.5% (2.3 logs) reduction was seen at the US EPA^13^, whereas Lab 2 reported a 99.72% (2.5 logs) and 97.77% (1.65 logs) reduction with the Clearify and the Aura devices, respectively. At the 15-minute sampling time, a 99.4% reduction (2.22 log) was reported at the US EPA, and a 99.9% (3 logs) reduction was reported for both the Clearify and the Aura devices tested at Lab 2.

Thus, testing conducted at Lab 1, Lab 2, and at the US EPA’s Office of Research and Development has consistently shown that GP can achieve up to a 3-log reduction in the concentration of airborne viable of the MS2 bacteriophage within 60 min of treatment and, in some cases, within 15 min of treatment.

Most of the conducted experiments are single experiments (i.e., each condition was not repeated by each lab). However, two separate laboratories conducted similar tests, and the results from their testing were consistent with each other as well as with the testing conducted by the US EPA. The totality of the evidence from the three sets of experiments indicates general agreement about the efficacy of GP against airborne viruses such as MS2 bacteriophage, which is often used as a surrogate for SARS-Cov-2.

### Toxicity of TEG – active ingredient of Grignard Pure

US EPA has concluded that TEG is of very low toxicity by the oral, dermal, and inhalation routes of exposure based on a review of available toxicology data^15^. The toxicology database is adequate to characterize the hazard of TEG, and no data gaps have been identified^15^. Further, the US EPA has not identified toxicological endpoints of concern for the active and inert uses of triethylene glycol. The US EPA has no risk concerns for TEG with respect to human exposure^15^. US EPA has also granted an exemption from the requirement of a tolerance for residues of antimicrobial pesticide ingredients for TEG (85 FR 69514) when used on or applied to food-contact surfaces in public places, including processing equipment^16^. TEG has also received “Generally Recognized As Safe” status for use as a food additive by the US FDA^17^.

TEG has been studied for repeat inhalation exposure effects in both rats and monkeys varying in duration from nine days to 13 months. In a nose-only exposure study, Sprague-Dawley rats were exposed to mean exposure concentrations of 102, 512, or 1036 mg/m^3^ of TEG for 6 hours a day for 9 consecutive days. In this study, no systemic adverse effects were seen at any level of exposure^18^. The investigators also concluded that “exposure to a respiratory aerosol is not acutely harmful, but may cause sensory irritant effects”^18^. Robertson et al (1947) conducted 3 different exposure studies with Monkeys. Browning of facial skin and crusting and damage to the skin of the ears occurred in 13 monkeys continuously exposed for 3 months or longer to an atmosphere containing about 4 mg TEG/m^3^ and described as ‘supersaturated.’ It was suggested that the bactericidal action of TEG may have promoted a parasitic infection which caused the skin damage^18^. Thirteen monkeys exposed continuously for 13 months to an atmosphere ‘supersaturated’ with TEG vapor (a concentration of 4 mg/m^3^) had slightly reduced weights ^18^. In a subsequent study, eight monkeys exposed to 2 – 3 mg/m^3^ for 10 months did not suffer skin effects, and no adverse effects were observed upon growth.

According to the European Chemicals Agency’s Classification and Labelling Inventory Database, TEG has not been classified as a human health hazard by the majority of the industry notifiers ^19^.

Nelson Laboratories, LLC (Salt Lake City, UT) developed a “Margin of Safety” (MOS) document for airborne TEG exposures based on Grignard Pure™ use levels for adult, child, infant, and neonatal populations utilizing the Tolerable Exposure limit (TE). Using standard toxicological exposure parameters such as tolerable intake, tolerable exposure, and published breathing rates and body weights for the various groups, MOS values were reported. By considering the maximum (worst-case) exposure to TEG (~ 1.0 mg/m^3^) from the use of the aerosolized GP product and the ISO 18562 default breathing rates for adults (20 m^3^/day), pediatrics (5 m^3^/day), infants (2 m^3^/day), and neonates (0.2 m^3^/day), MOS was calculated by dividing the TE by the worst exposure amount ^17^. The margin of safety for TEG under a worse-case exposure situation ranged from 2.0 for pediatric exposures to an average of 3.2 for adult men and women. For reference, an MOS value greater than 1 indicates a low toxicological hazard^17^. The report concludes that given these favorable MOS values, “acute, subacute/sub-chronic, and chronic toxicity, genotoxicity, and carcinogenicity from the exposure to TEG from the intended use of the product are not expected”^17^.

Another independent review of the safety TEG by TSG Consulting (Washington, DC) further concluded that the concentrations of airborne TEG from the use of the GP products (≤ 1.5 mg/m^3^) are more than 100 times less than the human equivalent concentration (~200 mg/m^3^) of the established limit dose for TEG in repeat-exposure animal inhalation toxicity studies (1,000 mg/m^3^)^20^.

In addition, the active ingredient of Grignard Pure™ – TEG has been utilized in lighting effects products that have been widely used for over two decades in theatrical, film, and TV productions, as well as at live events like concerts, sports, and worship services. The lighting effects product, which is used in a manner similar to GP, has exposed millions of people to concentrations of TEG between 5 – 10 mg/m^3^, often even at higher levels, and there have been no reported health issues associated with these exposures.

In summary, the results presented above by this study, as well as testing by the US EPA, show that the Grignard Pure™ (GP) product is able to inactivate over 99% of airborne virus particles within one minute of their introduction into an indoor space containing the product, and the inactivation reaches 2-3 logs within 60 minutes. In addition, there is a large body of scientific research indicating that the TEG levels at which GP is effective, e.g., 0.3 mg/m^3^ to 0.5 mg/m^3^ of GP, pose negligible health risks to humans.

The SARS-CoV-2 virus continues to mutate, and the newer emerging variants have proven to be more transmissible than earlier ones. As a result, despite vaccines, masking, and social distancing measures, the numbers of COVID-19 cases in the United States and globally have risen rapidly to levels higher than previously seen during the pandemic, putting immune-compromised and unvaccinated individuals at high risks of serious illness. A recent study by Lai et al. (2022) indicated that infected persons shed infectious SARS-CoV-2 aerosols even when fully vaccinated and boosted^21^. The evolutionary selection appears to have favored SARS-CoV-2 variants associated with higher viral aerosol shedding, requiring non-pharmaceutical interventions, especially indoor air hygiene (e.g., ventilation, filtration, and air disinfection) to mitigate COVID-19 transmission in vaccinated communities^21^. To minimize exposure to the virus and decrease the incidence of COVID-19 cases, it is critical to develop and utilize additional layers of protection.

One such layer could be the application of technological solutions to continuously inactivate the virus that is present or has been introduced into indoor space by infected individuals. Aerosolized TEG could be an important additional tool for lowering exposures to the SARS-CoV-2 virus in occupied and unoccupied indoor spaces. Compared to enhanced filtration and ventilation measures, GP provides a faster-acting mechanism to reduce airborne concentrations of the virus, as demonstrated by the testing described in this paper. Air change rates in typical occupied spaces range from 2-4 air changes per hour, resulting in a period of 90-180 min to complete 6 changes of room air. GP has been tested to provide a 99.5% reduction in airborne virus concentrations in a period of less than 10 min, accomplishing a 99% reduction in airborne concentrations 9 to 18 times faster than central ventilation, HEPA filtration, or UV treatment alone^22^. Therefore, it can be used as a continuous anti-virus air treatment either by itself or in conjunction with enhanced ventilation and air filtration measures. Maintaining a preset level of GP in the air of an indoor space would provide continuous protection to its occupants by inactivating a very high percentage of virus particles within minutes as they are newly introduced into the space. It could prove useful in spaces such as movie theaters, public transit vehicles, hotel rooms, offices, and other public spaces. Moreover, as an engineering control, everyone present in spaces where the product is used would receive its benefits, in contrast to vaccination, masking, and social distancing, all of which depend on individual choices for their success. Further testing of GP might even demonstrate its efficacy against other airborne pathogens such as the influenza virus.

The TEG-based antimicrobial air treatment product tested here shows high efficacy of viral inactivation and a favorable safety profile. As a result, it can be used to reduce exposure to the SARS-CoV-2 virus in indoor public spaces.

## Acknowledgments

We would like to extend our thanks to the support of Steve Winston and the Crisis Science Collaboration Group.

