## Supplemental Information for "The efficiency of Grignard Pure™ to inactivate airborne SARS-CoV-2 surrogate"

### **Measurement of Triethylene Glycol (TEG) and GP concentrations in the air**

We used the Hurricane Haze 1DX (Chauvet DJ, Sunrise, Florida) vaporizing device to generate a GP aerosol. Then we correlated the GP aerosol mass concentration, as measured by a handheld optical particle monitor DustTrak DRX (model 8534, TSI Inc., Shoreview, MN), with the total airborne TEG concentration (aerosol + vapor). The DRX is a multi-channel, battery-operated, data-logging, light-scattering laser photometer. It was calibrated by the manufacturer according to the ISO standard 12103-1 A1 using ultrafine Arizona Road Dust with a density of 2.65 g/cm^3^. The device was set to measure PM_10_ concentration. Before each test, the DRX was zeroed using the zero filter. Next, aerosol + vapor concentrations of TEG were measured using NIOSH Method 5523 (Pendergrass, 1996) with XAD-7 OVS sampling tubes (SKC, Inc., Eighty-Four, PA). The OVS tubes combine a sorbent and a filter inside a glass tube to trap aerosol particles and vapors simultaneously. Gillian GilAir3 (Sensidyne, St. Petersburg, FL) pumps provided a 1.2 L/min sampling flow rate for the tubes, and the samples were collected for 4.17 minutes for a total sample volume of 5 liters.

Grignard Pure Poster

36” x 36”

2ft

Fan 2

Vaporizing Machine

Fan air flow direction

DRX Aerosol Monitor

GilAir 3 Sampler

Fan 1

Fan air flow direction

Sampling Table

Figure S1. Schematic diagram of experimental set up for generation and measurement of TEG.

Room Door

Testing was conducted in a 1575 ft^3^ (44.5 m^3^) windowless chamber with dimensions 15 ft x 15 ft x 7 ft (Figure S1). The room was opened prior to and in between each experiment to facilitate the removal of GP aerosol and vapor. The removal of residual aerosol was verified utilizing the DRX aerosol monitor. Two small, portable Honeywell Home Table circulation fans were used at their highest setting during the entire experiment to disperse GP aerosol across the room. The Hurricane Haze 1DX (Chauvet DJ) was utilized to generate the 5 different airborne concentrations of GP (3.4 mg/m^3^ – 10.7 mg/m^3^). The vaporizing machine was placed 4 ft. above ground and 2 ft. from the Grignard Poster (Figure S1). The poster is used to visually assess the airborne GP level. The temperature in the room was between 20^o^C and 23^o^C, and the relative humidity was between 50 and 55 %.

The air pumps and sampling media were placed 5 ft. above ground, corresponding to the breathing zone for most people ^2^. Blank samples in duplicate were collected before the dispersion unit was turned on. The vaporizing dispersion unit was turned on during the experiments to reach a desired GP aerosol level, as measured by the DRX. Once GP aerosol concentration was stable, the TEG samples were collected using an OVS tube and GilAir3 sampling pumps.

All sample collection media and blanks were stored in a thermal bag with cooler packs and sent to an American Industrial Hygiene Association (AIHA)-accredited lab. All the samples were analyzed using NIOSH 5523 method with gas chromatography – flame ionization detection. The NIOSH method was extended to a validated limit of detection of 15.0 μg/L of TEG by the analytical lab (Analytics Corporation, Ashland, VA).

Since the DRX aerosol monitor was calibrated utilizing Arizona Road dust, which has a density of 2.7 g/cm^3^, a correction factor of 0.4 was applied to obtain the PM_10_ mass concentration of GP, which has a density of 1.08 g/cm^3^. In addition, the unadjusted GP aerosol mass concentration measured by the DRX was correlated with the total TEG airborne mass concentration (aerosol + vapor) measured by the OVS tubes and determined by the Analytics Corporation. The resulting linear regression yielded a slope coefficient of 0.23, the multiplication factor used to convert the real-time aerosol mass concentration measured by the DRX (not adjusted for density) into the total TEG concentration in the air. For example, an aerosol concentration of 30 mg/m^3^ measured by the DRX would correspond to 6.9 mg/m^3^ of TEG. The calibration curve for the relationship between aerosol concentration as measured by the DRX and total TEG levels is presented in Figure S2.

Total (aerosol + vapor) TEG concentration, mg/m3

| GP PM10 Aerosol Concentration, as measured by DRX, mg/m3 | TEG mg/m3 |
| --- | --- |
| 0.13 | 0 |
| 5.08 | 3 |
| 8.41 | 3 |
| 14.58 | 3.78 |
| 20.33 | 5.18 |
| 26.67 | 4.82 |

**Figure S2.** Calibration curve between Grignard Pure PM10 aerosol and total TEG concentration. The calibration factor, based on the slope of the curve, is 0.23.

**Table S1.** Correlation between GP PM10 aerosol concentrations as measured by DRX and the total TEG concentration (aerosol + vapor) at various Grignard Pure treatment levels.

| Grignard Pure Treatment Level | Average GP PM10 aerosol concentration, mg/m^3^ | Total TEG concentration, mg/m^3^ |
| --- | --- | --- |
| Non-Visible | 4.34 | < 3.00 |
| Very Light | 8.41 | < 3.00 |
| Light | 14.58 | 3.78 |
| Light Moderate | 20.33 | 5.18 |
| Moderate | 26.67 | 4.82 |

Glove Ports

Nebulizer

1

Nebulizer

2

Chamber Door

Particle Counters

Grignard Pure Vaporizing Device

Sampling Ports

Fan

**Figure S3**: Lab 1 Test Chamber Process Flow Diagram.

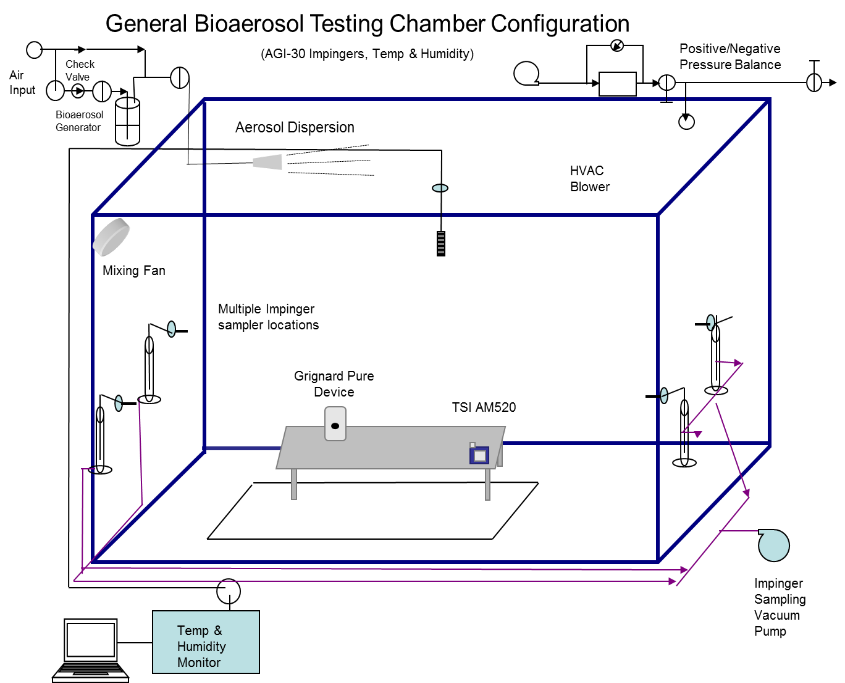

**Figure S4**: Lab 2 Test Chamber Process Flow Diagram.

**Table S2.** The concentration of viable airborne MS2 bacteriophage at different time points and the resulting reduction in viable airborne MS2 concentration when treated with a single four-second burst of Grignard Pure applied with the Nimbus, a handheld vaporizing device, compared to control concentrations at the same time points.

| **Experiments** | **Start of sampling relative to introducing GP, min** | **Viable airborne MS2 concentration, PFU/m^3^** | **Gross log reduction in viable airborne MS2 concentration compared to t = -10 min** | **Percent gross reduction in viable airborne MS2 concentration compared to t = -10 min** | **Net log reduction in viable airborne MS2 concentration due to air treatment by GP** | **Percent net reduction in viable airborne MS2 concentration due to air treatment by GP** |
| --- | --- | --- | --- | --- | --- | --- |
| Control experiment, Lab 1 | -10 | 3.42E+08 | N/A | N/A | N/A | |
|  | 0.5 | 2.43E+08 | 0.15 | 28.947 |  |  |
|  | 15 | 1.23E+08 | 0.44 | 64.035 |  |  |
|  | 60 | 2.94E+07 | 1.07 | 91.404 |  |  |
| Treatment experiment, Lab 1 | -10 | 2.57E+08 | N/A | N/A | N/A | |
|  | 0.5 | 3.91E+05 | 2.82 | 99.848 | 2.67 | 99.786 |
|  | 15 | 1.19E+05 | 3.33 | 99.954 | 2.89 | 99.871 |
|  | 60 | 7.96E+04 | 3.51 | 99.969 | 2.44 | 99.640 |
| Control experiment, Lab 2 | -10 | 6.67E+10 | N/A | N/A | N/A | |
|  | 0.5 | 4.59E+10 | 0.16 | 31.184 |  |  |
|  | 15 | 1.49E+10 | 0.65 | 77.661 |  |  |
|  | 60 | 3.87E+09 | 1.24 | 94.198 |  |  |
| Treatment experiment, Lab 2 | -10 | 4.27E+10 | N/A | N/A | N/A | |
|  | 0.5 | 6.16E+07 | 2.84 | 99.856 | 2.68 | 99.790 |
|  | 15 | 6.24E+06 | 3.84 | 99.985 | 3.18 | 99.935 |
|  | 60 | 2.64E+06 | 4.21 | 99.994 | 2.97 | 99.893 |

Table S3. The concentration of viable airborne MS2 bacteriophage at different time points and the resulting reduction in airborne viable MS2 bacteriophage concentration when treated with time-controlled release of different concentrations of GP and the resulting TEG. The Grignard Pure was released in the time-controlled method using the Amhaze, a stand-alone vaporizing device.

| **Experiments** | **Start of sampling relative to introducing GP, min** | **Airborne MS2 concentration, PFU/m^3^** | **Gross log reduction in airborne PFU concentration compared to t= -10 min** | **Gross percent reduction in airborne PFU concentration compared to t= -10 min** | **Net log reduction in airborne PFU concentration due to air treatment by GP** | **Percent net reduction in airborne PFU concentration due to air treatment by GP** | |
| --- | --- | --- | --- | --- | --- | --- | --- |
| Control Experiment 1 | -10 | 1.25E+09 | 0.00 | 0.00 | N/A | | |
|  | 0.5 | 1.20E+09 | 0.02 | 4.270 |  |  |  |
|  | 15 | 9.55E+08 | 0.12 | 23.600 |  |  |  |
|  | 60 | 5.43E+08 | 0.36 | 56.590 |  |  |  |
| Treatment Experiment 1 (TEG - 0.186mg/m³) | -10 | 1.77E+09 | 0.00 | 0.000 | 0.00 | 0.00 | |
|  | 0.5 | 1.86E+07 | 1.98 | 98.949 | 1.96 | 98.905 | |
|  | 15 | 8.44E+06 | 2.32 | 99.523 | 2.20 | 99.376 | |
|  | 60 | 1.37E+06 | 3.11 | 99.923 | 2.75 | 99.822 | |
| Control Experiment 2 | -10 | 7.79E+08 | 0.00 | 0.00 | N/A | | |
|  | 0.5 | 7.26E+08 | 0.03 | 6.804 |  |  |  |
|  | 15 | 3.52E+08 | 0.35 | 54.814 |  |  |  |
|  | 60 | 4.64E+07 | 1.23 | 94.044 |  |  |  |
| Treatment Experiment 2 (TEG - 0.287mg/m³) | -10 | 1.45E+08 | 0.00 | 0.00 | 0.00 | 0.00 | |
|  | 0.5 | 2.59E+06 | 1.75 | 98.214 | 1.72 | 98.083 | |
|  | 15 | 5.51E+04 | 3.42 | 99.962 | 3.08 | 99.916 | |
|  | 60 | 1.33E+04 | 4.04 | 99.991 | 2.81 | 99.846 | |
| Control Experiment 3 | -10 | 3.80E+08 | 0.00 | 0.00 | N/A | | |
|  | 0.5 | 4.13E+08 | -0.04 | -8.648 |  |  |  |
|  | 15 | 1.73E+08 | 0.34 | 54.474 |  |  |  |
|  | 60 | 4.25E+07 | 0.95 | 88.816 |  |  |  |
| Treatment Experiment 1 (TEG 0.235mg/m³) | -10 | 1.35E+08 | 0 | 0 | 0 | 0 | |
|  | 0.5 | 5.67E+05 | 2.38 | 99.581 | 2.38 | 99.58 | |
|  | 15 | 1.05E+05 | 3.11 | 99.923 | 2.77 | 99.830 | |
|  | 60 | 9.81E+03 | 4.14 | 99.993 | 3.19 | 99.935 | |
| Treatment Experiment 2 (TEG 0.235mg/m³) | -10 | 4.51E+07 | 0.00 | 0.00 | 0 | 0 | |
|  | 0.5 | 2.12E+05 | 2.33 | 99.529 | 2.33 | 99.53 | |
|  | 15 | 1.02E+05 | 2.65 | 99.775 | 2.31 | 99.505 | |
|  | 60 | 1.23E+04 | 3.57 | 99.973 | 2.61 | 99.757 | |
| Treatment Experiment 2 (TEG 0.235mg/m³) | -10 | 1.98E+07 | 0.00 | 0.00 | 0 | 0 | |
|  | 0.5 | 1.01E+05 | 2.29 | 99.491 | 2.29 | 99.49 | |
|  | 15 | 6.10E+04 | 2.51 | 99.692 | 1.56 | 97.248 | |
| Control Experiment 4 | -10 | 4.62E+08 | 0.00 | 0.00 | N/A | | |
|  | 0.5 | 3.29E+08 | 0.15 | 28.788 |  |  |  |
|  | 15 | 1.90E+08 | 0.39 | 58.874 |  |  |  |
|  | 60 | 4.69E+07 | 0.99 | 89.848 |  |  |  |
| Treatment Experiment 4(TEG - 0.063g/m³) | -10 | 8.99E+07 | 0.00 | 0.00 | 0.00 | | 0.00 |
|  | 0.5 | 4.88E+06 | 1.27 | 94.572 | 1.12 | | 92.390 |
|  | 15 | 2.69E+05 | 2.52 | 99.701 | 2.14 | | 99.270 |
|  | 60 | 5.98E+04 | 3.18 | 99.933 | 2.18 | | 99.340 |

Table S4. The concentration of viable airborne MS2 bacteriophage at different time points and the resulting reduction in airborne viable MS2 bacteriophage concentration when treated with time-controlled release of different concentrations of GP and the resulting TEG. The GP was aerosolized using the Clearify, a handheld vaporizing device, and the Aura, a stand-alone nebulizing device at Lab 2.

| **Experiments** | **Start of sampling time, t*, min** | **Airborne MS2 concentration, PFU/m^3^** | **Gross log reduction in airborne PFU concentration compared to t*= -10 min** | **Percent gross reduction in airborne PFU concentration compared to t* = -10 min** | **Net log reduction in airborne PFU concentration due to air treatment by GP** | **Percent net reduction in airborne PFU concentration due to air treatment by GP** |
| --- | --- | --- | --- | --- | --- | --- |
| Control experiment, Lab 2 | -10 | 1.87E+10 | 0.00 | 0.000 | N/A | |
|  | 15 | 7.31E+09 | 0.41 | 60.909 |  |  |
|  | 30 | 4.19E+09 | 0.65 | 77.594 |  |  |
|  | 60 | 2.00E+09 | 0.97 | 89.305 |  |  |
|  | 90 | 1.47E+09 | 1.10 | 92.139 |  |  |
|  | 120 | 5.55E+08 | 1.53 | 97.032 |  |  |
| Treatment Experiment, Lab 2 (Clearify device) TEG - 0.092mg/m³ | -10 | 5.09E+07 | 2.56 | 99.7277 | 2.56 | 99.7277 |
|  | 15 | 4.29E+06 | 3.64 | 99.9770 | 3.23 | 99.9413 |
|  | 20 | 1.89E+06 | 3.99 | 99.9899 | 3.35 | 99.9549 |
|  | 60 | 1.31E+06 | 4.15 | 99.9930 | 3.18 | 99.9345 |
|  | 90 | 7.63E+05 | 4.39 | 99.9959 | 3.28 | 99.9481 |
|  | 120 | 4.59E+05 | 4.61 | 99.9975 | 3.08 | 99.9173 |
| Treatment Experiment, Lab 2 (Aura device) TEG - 0.025mg/m³ | -10 | 4.16E+08 | 1.65 | 97.7714 | 1.65 | 97.7754 |
|  | 15 | 1.81E+07 | 3.01 | 99.9029 | 2.61 | 99.7524 |
|  | 20 | 4.37E+06 | 3.63 | 99.9766 | 2.98 | 99.8957 |
|  | 60 | 1.81E+06 | 4.01 | 99.9903 | 3.04 | 99.9095 |
|  | 90 | 7.15E+05 | 4.42 | 99.9962 | 3.31 | 99.9514 |
|  | 120 | 2.85E+05 | 4.82 | 99.9985 | 3.29 | 99.9486 |
